# Quantitative Late Quaternary climate reconstruction from plant macrofossil proxy in Western North America

**DOI:** 10.1101/340208

**Authors:** Robert S. Harbert, Kevin C. Nixon

## Abstract

The Late Quaternary packrat (*Neotoma* spp.) midden plant macrofossil record in western North America is an exceptional record of biotic change that provides strong evidence of past climate. In this study we generate quantitative estimates of climate from plant community composition of more than 600 individual paleomiddens over the past 50,000 years. This is the first large-scale application of CRACLE, a quantitative climate inference method that uses plant community composition as a climatic proxy under and individualistic concept of plant community assembly. The results are spatiotemporally specific estimates of temperature, precipitation, available moisture, and seasonal patterns that are consistent with well understood global climate patterns, but provide previously unavailable detail and precision of the regional paleoclimate in western North America. Rapid warming is estimated at the Pleistocene-Holocene transition, at a conservative estimate of ca. 1°C per millennium. Previously projected future temperature increases suggest a rate of increase of more than 2°C over the next century, an astonishing 10X the rate experienced at any point during the past 50,000 years in Western North America. These analyses form a baseline demonstration of how the growing paleoecological record of packrat midden plant macrofossils is able to provide quantitative estimates of paleoclimate that aid in understanding the complexities of, and biotic responses to the regional climate system.

## INTRODUCTION

Past mass extinction and spatial shifts in biodiversity due to climate change can provide critical data to understand the potential future effects of climate change on global and regional ecosystems. Geologically rapid temperature and precipitation fluctuations during the Quaternary glaciation cycles (1,2) have been used as analogs for potential biotic change during projected near-future climate change (e.g., 3; 4). These past changes are believed to have been largely driven by predictable and periodic variation in the earth’s orbit relative to the sun (5,6) effecting changes in relative insolation (7). The global glacial record is well-preserved and continuous over thousands of years in both the arctic and antarctic ice-sheets, and preserves representative atmospheric gasses that are indicators of past climate (8,9,10). However, North American midlatitude quantitative climate records are either less continuous or over a short time scale (11) and primarily limited to ocean and lake sediment analyses including isotopes and diatoms (e.g., 12,13,14), groundwater stable isotope analyses (e.g., 15,16), and scattered biological proxies (e.g., 17,18,19,20). Late Quaternary plant macrofossils from packrat paleomiddens in western North America as presented here therefore represent a significant, independent, and novel source of data for large-scale mid-latitude paleoclimate inference for western North America.

### North American Packrat Paleomiddens

North American packrat (*Neotoma* spp.) paleomiddens (fossilized nests) have been a rich source of data for Late Quaternary paleoecological estimates (21). Packrats rather indiscriminately collect both vegetative and reproductive plant material from the local environment to build nests and to establish caches of food. These collections, including leaf and stem fragments, fruits, seeds, and pollen blown into the midden matrix (22) typically form a representative sample (60-90% of taxa) of the local plant diversity within 30-50m of a midden (23,24). Because of the excellent preservation of the materials in dry, protected cavities and under ledges, the fossils can typically be identified morphologically as belonging to living species from western North America. Paleomidden analysis has long been an important approach to documenting changes in vegetation, population migration and colonization, and morphological and physiological characterization of vegetation of the Late Quaternary as far back as 50,000 years before present (*e.g*., 25, 26, 27, 28, 29, 30, 31, 32, 33, 34, 35, 36, 37, 38, 39). Middens are often chronologically layered and these layers can be dated with radioisotope methods to provide a sequential history of vegetation composition. The packrat paleomidden fossil record shows remarkably complete biogeographic histories that facilitate the study of past distributional shifts due to local extinction and dispersal processes (e.g. 39, 40), critical information for understanding how plants respond in natural environments to periods of rapid climate change.

### Paleoclimate estimation from Vegetation

Climate Reconstruction Analysis using Coexistence Likelihood Estimation (*CRACLE*) — The principle that plant species distributions are intimately correlated with climate across broad spatial and temporal scales is universally accepted (e.g., 41). This principle is the basis for a class of climate estimation techniques that use climatic niche overlap between coexisting species to estimate site-specific climate parameters (Coexistence Approach: (42); Mutual Climatic Range: (43,44,45)). In this study, climate parameters are estimated using a maximum likelihood statistical framework for the estimation of climate from vegetation via CRACLE, Climate Reconstruction Analysis using Coexistence Likelihood Estimation (46). CRACLE is a relatively new, but robust method that leverages large online databases of primary biodiversity data such as the Global Biodiversity Information Facility — GBIF (www.gbif.org) — and has been shown to provide more accurate estimates of climate than related methods (46). Previously, Mutual Climatic Range methods have been applied to the study of Late Quaternary climate for only a few packrat paleomidden samples (44,45), but a comprehensive regional study has not been completed. This study is the first to apply a taxonomic coexistence climate estimation method to the entire body of data available for Late Quaternary packrat paleomiddens in western North America. It is also the first large-scale application of the CRACLE implementation to paleoclimate modeling (47).

## RESULTS AND DISCUSSION

### CRACLE Modern Validation Experiment

To validate CRACLE as a sensitive and robust method for estimating climate across the Late Quaternary of western North America a modern set of more than 3,493 vegetation plot surveys covering North America west of 100°W and between 25°N and 50°N (mobilized via the Botanical Information and Ecology Network — BIEN — http://bien.nceas.ucsb.edu/bien/) database were analyzed using the same CRACLE methodology as for the packrat midden fossil communities (All code available at: https://github.com/rsh249/cracle_examples/tree/master/BIEN).

These modern sites cover between -3°C to 23°C degrees of mean annual temperature and 100-1,100mm of annual rainfall, a latitudinal gradient of ~20 degrees, and altitudes between 170 and 4000 meters. This dataset should well represent the scope of climatic space for the Late Quaternary of this part of western North America.

Temperature variables (e.g. Mean Annual Temperature — MAT) were estimated with high correlation (ρ = 0.93) and low error rates (median absolute anomaly = 1.3°C — Fig. S1; Supplementary Table S1) relative to the WorldClim estimates for those sites. Estimation of precipitation parameters (mean annual precipitation and annual water balance) exhibited greater variation in results, but with median anomalies of 79mm for mean annual precipitation and 96mm for annual water balance (Table S1). Seasonal estimation, represented in this study by winter length (the number of months with mean temperatures below 5°C), performs well with median anomaly of only 0.05 months, or about 1.5 days. The performance in this regional dataset is comparable to the global set of test sites originally published for CRACLE (46), but from a dataset almost 25 times larger. These results show that CRACLE is a robust tool in for this regional flora and the climate space relevant to the packrat midden macrofossil record of the Late Quaternary.

### Paleoclimate Overview

The climate record inferred by CRACLE analysis of the 646 dated western North American packrat paleomidden samples is consistent with patterns of northern hemisphere temperature changes over the last ~50,000 years based on other methods (8,9,10). Coldest temperatures with anomalies of −2 to −4°C were estimated between 22kya and 15kya, during the last glacial maximum (LGM) in Western North America. The late Pleistocene temperature minima were followed by a general warming trend that continued until after the onset of the Holocene, peaking with positive anomalies of +1.5°C above modern values consistent with the Holocene Thermal Optimum (48) at 7-8kya (Fig. 1). Atmospheric CO2 recorded in the Antarctic ice cores (Fig. 1E) (12) increases ahead of mean annual temperature increases in the CRACLE inferred record (Fig. 1A) supporting previous observations of global climate change at the end of the Pleistocene (49). Although showing dramatic extremes (~6°C total increase in temperature), the magnitude of CRACLE-packrat estimated mean temperature shifts at these latitudes is much less than is observed in the reliable ice-core proxies of Greenland (8,9,10). This is consistent with a scientific consensus that global temperature change and ecosystem response is typically greater in polar regions during historic warming events (48, 50).

### Rate of Climate Change

The maximum rate of mean annual temperature change estimated in this study (~6°C/6 to 8 thousand years = 0.075 - 0.1°C/100 years; Fig. 1) is a slower rate than current projections of near-future warming scenarios for western North America. The projected rate of warming across the western United States through the year 2100 is projected to be between 1.5 and 4°C in all IPCC Representative Concentration Pathways, including the stringent mitigation scenario—RCP 2.6— that is, at this point, very unlikely to be attained (51). In the Southwestern United States, recent historical records suggest increasing temperatures are now occurring at a rate of ~3-4°C/100years, a rate consistent with model projections for the next century (52). Based on these conservative estimates contrasted against CRACLE paleomidden estimates, projected warming in the next century is likely to occur at a rate 10-20X greater than any millennial average in the past 50,000 years (52). This disconnect highlights the need for studies that infer potential future biotic change (e.g., 53) and migration rates (54) to consider the unparalleled effects of very rapid periods of climatic change rather than just the total magnitude of projected change (55).

### Pleistocene Pluvial Lakes

One of the most striking geologic features of the late Pleistocene is the occurrence of large lake systems in the Great Basin during the Last Glacial Maximum (LGM). Estimates of mean annual precipitation (MAP) and water balance (MAP - potential evaporation) for the LGM are consistent with a wet-glacial hypothesis (56), with additional rainfall of 50-100mm/year over modern values (Fig. 2C). Driven by cooler temperatures and reduced evaporation, relative water balance (i.e., subtracting estimated evaporation) was greater than modern values by +200mm/year to +400mm/year from 22kya to at least 15kya (Fig. 2D). During the Bolling-Allerod interstadial (a pronounced Northern Hemisphere warming episode from 14.7kya to 12.7kya), the wetter regime across the region started to collapse. At this time, CRACLE paleomidden analyses show rapidly diminishing rainfall from 15kya towards the modern regime by ~12kya. This decrease in rainfall is followed closely, though with less abrupt changes, by water balance decreases as temperatures increased towards the HTO (Fig. 2A, 2B). Interestingly, CRACLE-estimated winter length remained longer than the modern by about one month until around 9kya (Fig. 2E), despite rising annual (Fig. 1A) and seasonal extreme temperatures (Fig. 2A, 2B).

The rise of the Pleistocene pluvial lakes in the Great Basin have been attributed to an influx of moisture due to increased rainfall and/or limited evaporation by the full glacial cold (14,16,56,57,58). Our estimated changes in water balance during the LGM (22 to 15kya) show a positive linear relationship with latitude — larger positive anomalies in water balance to the north — (Fig. 3). This is consistent with other contemporary models that show greater rates of temperature increase in northern, more polar latitudes than in southern or more tropical latitudes (48, 50). Predicted differences in water balance were used to adjust the modern climate model grid using a linear model of latitude vs. estimated water balance change (Fig. 3) for the samples from the LGM maximum positive anomalies and mid-holocene maximum negative anomalies. During the LGM these models produce anomalies of up to +500 mm/year of water balance, with greater values in the north, pushing positive water balance regimes to the south and down-slope.

### The Holocene: Implications for future drought scenarios

Based on CRACLE estimates, much of the Holocene is characterized as warmer by 1-2°C, and dryer than the present (Fig. 1A). Our reconstructions of precipitation and water balance suggest that the Holocene after ~8kya was as dry as, or dryer than, the current climate throughout the western U.S. However, it is important to note that current near-future climate projections indicate we will exceed this worst-case historical scenario, and perhaps more importantly, at a much quicker rate.

Temperature maxima in the Holocene are estimated to occur at 7-8kya and 2-4kya for all temperature parameters (Fig. 2A,B). In turn, temperature increases appear to drive estimated water balance minima at those same time periods (Fig. 2D). This result is in contrast to an initial warming followed by a steady downward trend in MAT between the HTO and the present reported from multiproxy climate reconstruction syntheses (15). The CRACLE estimated warm periods and the pattern of associated expansion of drought regimes (Fig. 3) are reasonable analogs for future warm climates in arid and semi-arid regions of western North America.

CRACLE estimates of drought (negative water balance) suggest that parts of the southern half of our study area, including Southern California, Arizona, and northern Mexico, may see shifts from positive to negative water balance regimes if the projected warming by year 2100 is realized (Fig. 3).

Estimation of Holocene water balance also reveals that these warm and dry climate periods had less impact to the north of the study area. In particular, the northern Sierra Nevada range and coastal regions of northern California, Oregon, and Washington show little change in the overall wet climate through the Holocene. This moisture regime stability may imply that these regions are historical, and potential future refugia. It is interesting that these regions (such as the Siskyou Range) are considered refugia for many plants that were more widespread in the past (59,60,61).

### Holocene Temperature Variability

All estimates of temperature presented here (mean annual, minimum, and maximum) show marked variability during the Holocene (Fig. 1A, 2A,B) including a local temperature minimum around 4-5kya. The dominant global temperature trend of the Holocene has been temperature stability with the exception of the last 2000 years where anomalous cooling has been inferred (15; 62). However, climate variability at local and regional scales may be missed in such broad syntheses. CRACLE analysis reconstructs an overall warm Holocene with a cooling trend over the last 2,000 years of the Holocene consistent with these global trends (15).

The CRACLE temperature estimates show first a ~2C cooling trend between 5-7kya followed by an equal warming trend between 3-5kya (Fig. 1A, 2A,B). Model data suggest a 2-4C maximum temperature anomaly during the early Holocene (48) in this region, an inference corroborated by the early warm period in the CRACLE estimates (Fig. 1A). Sea surface temperature (SST) along the Northern California coast may have been cooler than present during the mid-Holocene (13). Diatom records from the northern Gulf of Mexico suggest an amplified North American Monsoon (NAM) resulting in a wet summer pattern during the mid-Holocene as well (63). Cooler SST (13) and increased summer rainfall (63) may help explain the temperature minimum at 5kya in our results as regional climatic response.

### Millennial Climate Variation and Plant Biodiversity Dynamics

Our CRACLE estimates for the end of the Pleistocene show that mean annual temperature increased from the LGM minimum prior to decreases in precipitation near the Pleistocene-Holocene transition (Fig. 1). Surprisingly, this period is also identified as having a significant millennial-scale increase in average within-midden plant diversity—temporal α-diversity— (Supplementary Fig. S2). Plant diversity in the arid regions of the Southwest is lowest in dense, high elevation forest, low elevation grasslands, and desert scrub, but highest in open, dry woodlands due to a mixing of both woodland and grassland/desert elements in this zone (64). At the end of the Pleistocene a warming trend with moderate rainfall that persisted across large portions of this region could have driven an expansion of a semiarid climate with open woodland vegetation types. Thus, increases in open woodland vegetation with relatively higher species diversity is one potential mechanism for the observed increase in within-midden diversity in this period (Supplementary Fig. S2).

## CONCLUSIONS

The Late Quaternary plant macrofossil record preserved in packrat paleomiddens provides exceptionally strong evidence to estimate a detailed and well-supported timeline of climatic change across western North America. This is the first application of the CRACLE approach to Pleistocene paleoclimate estimation and the first large-scale quantitative reconstruction of climate in this region using plant macrofossils and demonstrates the potential application of CRACLE to other paleoecological records and databases (e.g., the Neotoma Paleoecology Database — 65). The 50,000-year climate profile estimated using CRACLE includes many well-understood features of global climate history found with other methods and in other regions, validating the overall approach and its application in this region. Perhaps more importantly, this reconstruction quantifies changes in numerous climate variables not typically accessible through other proxy methods (e.g., water balance and winter length). The climate profile also sheds light on key events of the geologic and fossil record including the development of large pluvial lakes in the Great Basin during the terminal Pleistocene glacial cycle. The rapid temperature increase, decrease in precipitation, reduced water balance and implied greater drought marking the Pleistocene-Holocene transition is evidence of climate change that elsewhere has been implicated in coordinated extinctions of Pleistocene megafauna in North America (66,67). However, future projected anthropogenic climate change may occur at a rate of ca. 1-4°C per century (51,52), at least one order of magnitude greater than the millennial rate of about 0.1°C per century estimated here for the Pleistocene-Holocene transition. The rate estimated in this study for the Pleistocene-Holocene transition is less than the ca. 0.5°C per century for polar regions for the same time period inferred from the GISP2 cores (8,9,10), highlighting the utility of the continuous mid-latitude climate reconstructions made by CRACLE using the packrat midden macrofossil system (or other similar biological data sources) for aiding the quantification of the latitudinal climate change differential. Climate shifts in the past have resulted mostly in rearrangements of the standing biodiversity of the regional flora of western North America rather than extinction. The generally modest rates of climatic change may have allowed for processes of migration and dispersal to track environmental shifts over long periods. Given the rapid changes projected for the next century, there appears to be no broad analog in the Late Quaternary of western North America for the rapidity of projected climate change that the world faces in the next century.

## METHODS

### CRACLE

Estimation of climate was computed based on taxon coexistence and modern distributions via the Climate Reconstruction Analysis Using Coexistence Likelihood Estimation (CRACLE) protocol (46). For each taxon in a community, CRACLE calculates a probability function representing the conditional probability of a climate parameter given the presence of that taxon. The joint likelihood function for all co-occurring species is then calculated as the sum-log-likelihoods of these individual taxon functions. The maximum of the joint likelihood curve is taken to be the most probable climate value given the association of species and their individual association with climate. Analyses in this study were done using the ‘*vegdistmod*’ R package for CRACLE and related methods (https://github.com/rsh249/vegdistmod). The methods used here differ from those originally published (46) in two ways. First, a weighted Kernel Density Estimation is used to approximate the conditional probability based on a background data sample. Second, standard bandwidths are calculated using Silverman’s Rule (68).

CRACLE estimates fossil taxon climate tolerance probabilities from modern primary biodiversity distribution data using a Smallest Inclusive Group (SIG) approach to identify proxy distributions for extinct or un-identifiable fossil taxa. Here, a SIG is defined as the smallest taxonomic group or clade that a fossil can be reasonably placed in using morphological characteristics. The taxonomic classifications from the USGS North American Packrat Midden Database (https://geochange.er.usgs.gov/midden/) are used as SIGs when the classification is made to the genus or species level. One assumption of this approach is that fossils only identified to family level will lack narrow climate signals due to the use of overly broad geographic distributions and likely could be placed into smaller groups with deeper study. It is expected that taxa in the packrat midden samples represent modern taxa as extinctions during this time are rare but rearrangement of species into novel communities was common (39,69). When generic classifications are used in the CRACLE framework (46) the distribution for the entire genus is substituted as a surrogate for the unknown, but extant, species distribution under the assumption of physiological canalization related to climate niche (70,71,72). Using modern species/genus distribution data for fossils that clearly fit within these groups avoids issues of niche shifts and extinction associated with applying this approach to older fossil assemblages that may lack taxa identifiable as equivalent to modern species (73).

### USGS-NOAA Packrat Midden Database

The work of countless researchers since the discovery of Pleistocene age *Neotoma* middens (25) to locate, catalog, and identify fossil plant remains in the middens of *Neotoma* spp. packrats is now largely curated by the United States Geological Survey in the form of the USGS-NOAA North American Packrat Midden Database (available online: http://geochange.er.usgs.gov/midden/search.html). The USGS North American Packrat Midden Database (https://geochange.er.usgs.gov/midden/) includes more than 3,000 midden samples, with species identifications, georeferenced localities, and radiocarbon estimated ages of plant material. Because of the historical nature of the dataset, some records lack some information. Sites may have a single age, or more often multiple, identified layers from different depositional times that can be sampled separately. For our study, the data set was filtered to remove samples without geo-coordinates, with poor coordinate precision, and those with fewer than 10 taxa identified to genus or species. The culled data set presented here includes 646 samples (Supplementary material Appendix 2) distributed from the deserts of northern Mexico to the northern Great Basin, eastern California, southern Oregon and Idaho (Supplementary Fig. S3). Rarity of the most ancient middens and radiocarbon dating uncertainties restrict usable samples for our study to the last 50,000 years with much greater representation of the last 10,000 years (Supplementary Fig. S2). On average, there were about 15-20 taxa identified per sample (Supplementary Fig. S2; **Distribution data used will be submitted to www.datadryad.org prior to publication**), a level of diversity that has been shown to be sufficient for accurate and precise CRACLE models (46).

### Midden Age Chronology and Paleoclimate Timeline --

Radiocarbon dates for the packrat middens were calibrated using the Northern Hemisphere IntCal13 calibration curve for terrestrial samples (74), and the calibration tools available in the “Bchron” software library (75) in R (76). Age calibration model means and standard deviations were converted to 95% confidence intervals for each site to represent temporal uncertainty in climatic reconstruction. Reconstructions average across all overlapping midden date ranges at each time bin.

### Modern climate model and parameters

Climate data are extracted from the downscaled 2.5-arcminute resolution (~0.041667 degrees) WorldClim (Version 1.4) model grid (77). WorldClim is a high-resolution continuous grid of interpolated climate data for the world’s land areas derived from a global set of >40,000 weather stations.

An advantage of the CRACLE method for inferring paleoclimate is that the general protocol is flexible and can be applied to a wide variety of climatic parameters. For this study quantitative reconstructions of mean annual temperature (MAT), maximum temperature, minimum temperature, mean precipitation, mean water balance (potential evapotranspiration + mean precipitation), and winter length (number of months with mean temperature < 5°C) are focused on as these give a broad view of climate change and seasonal variability (Supplementary material Appendix 1).

To better capture nuances of climate related to drought, the potential evapotranspiration (PET) and water balance (precipitation – PET) was calculated using monthly values for temperature and precipitation from the WorldClim model (77) and in R (using code from: https://github.com/rsh249/raster_PET.git). The Thornthwaite Equation (78,79) was chosen as a suitable model of potential evapotranspiration that relied on data available at the scale of WorldClim including: 1) monthly average temperatures in degrees Celcius, 2) mean monthly day-length in hours (calculated from the latitude), and 3) the number of days in each month.

### Primary Biodiversity Data

Point-locality distribution data were downloaded from the Global Biodiversity Information Facility API service (http://api.gbif.org/v1/). For each taxon (genus or species) queried, localities were removed if any of the following criteria were met: 1) Decimal longitude or latitude was reported as an integer or to only a single decimal place, implying poor precision. 2) The basis of record was listed as “FOSSIL_SPECIMEN”. 3) The locality description had any word matching in-whole or with up to two mismatches “botanic”, “garden”, or “cultivated” to remove observations from cultivated settings. To reduce the effect of spatially biased collection patterns all data were thinned by species to a 20 arcminute grid (the 2.5 arcminute WorldClim grid aggregated by a factor of eight). Taxa with fewer than 5 records after filtering were excluded as incomplete data. The final occurrence dataset consisted of 368,702 unique records for the 341 genera and 533 species observed in the 646 studied midden samples (**Exact distribution data used will be submitted to www.datadryad.org or similar public repository prior to publication**).

### Regional CRACLE Validation Testing

To test whether CRACLE is adequate for capturing climate information at a regional scale comparable to the total climate space for the Late Quaternary of western North America the model was applied to vegetation surveys from the United States, Canada, and Mexico mobilized by the Botanical Information and Ecology Network (BIEN, http://bien.nceas.ucsb.edu/bien/) via the R library ‘rbien’ (80). For each survey primary biodiversity data from GBIF was download as described above and all the same filtering and analytical steps were applied as with the USGS Packrat Midden Database. In the final distribution data there were 5,563 species and 915,414 records. One additional filter was applied for the modern sites which was to remove surveys with reported elevation greater than 200 meters above or below the estimated elevation used in the WorldClim model (77; http://srtm.csi.cgiar.org/). Elevation filtering will reduce calculated anomalies by removing surveys that are either poorly georeferenced or were collected on a very strong elevation gradient that is not captured well in the WorldClim climate interpolation. All code and data to reproduce this validation test is publicly available (https://github.com/rsh249/cracle_examples/tree/master/BIEN).

**Figure 1.**
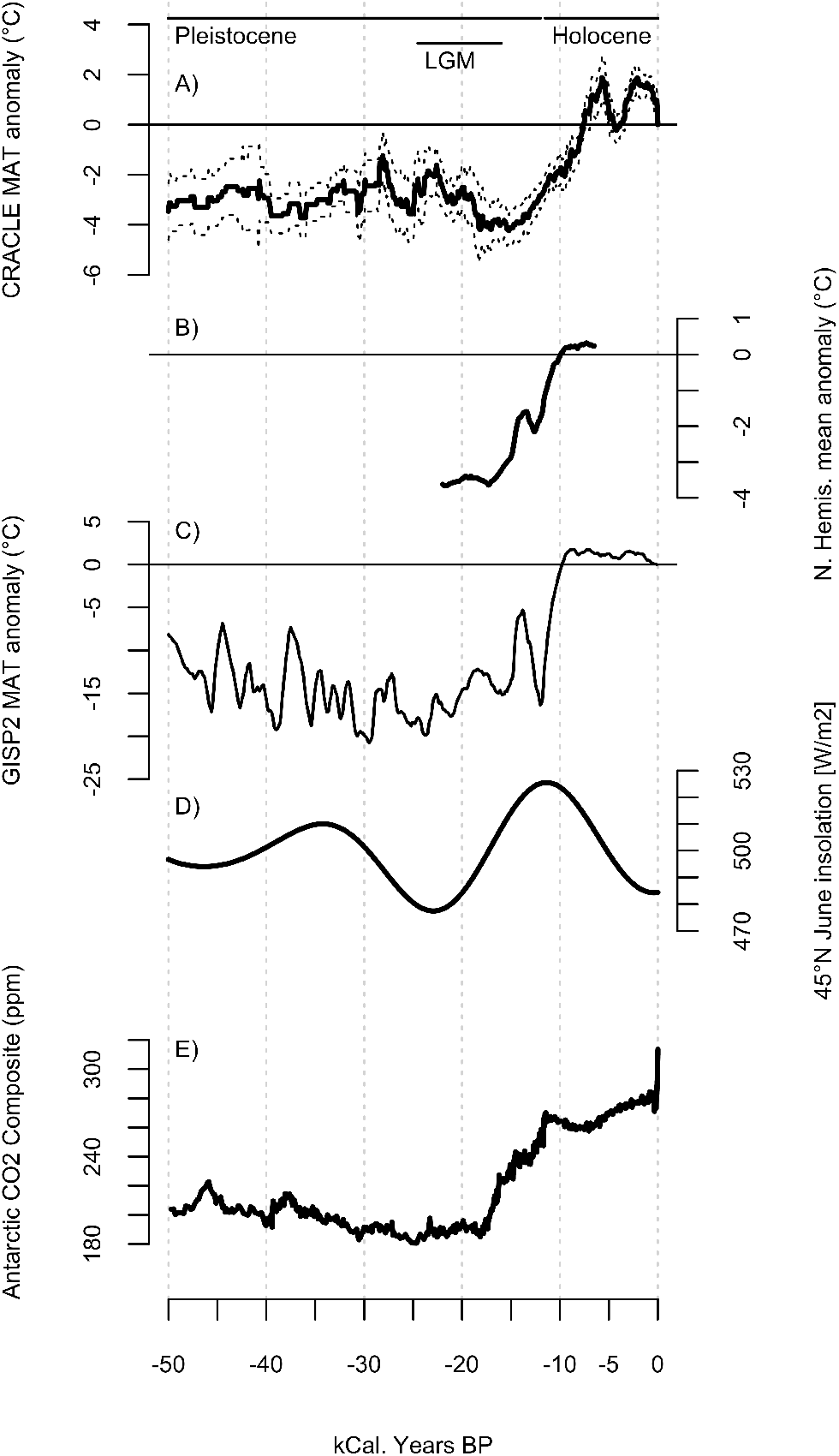
50,000 years of CRACLE Mean Annual Temperature record, Northern Hemisphere Ice-Core estimated temperature data, peak summer insolation at 45N, and atmosphere CO2 concentration. A) Sliding window 1000 year moving average Mean Annual Temperature (MAT) difference estimated by CRACLE shown as the mean deviation +/− 2 standard error (dashed lines) from modern average for the 646 study sites. B) Northern Hemisphere multiproxy temperature anomaly stack (48). C) Greenland air temperature sliding window 1000 year moving average estimated anomaly (Data from the Greenland Ice Sheet Project 2, ftp://ftp.ncdc.noaa.gov/pub/data/paleo/icecore/greenland/summit/gisp2/isotopes/gisp2_temp_accum_alley2000.txt 8,9,10). D) Mean daily summer (June) insolation at 45N (kJ/m2) estimated following the geometric method (3) using the ‘palinsol’ R library (81). E) Mean CO2 concentration (ppm) estimated from a composite of Antarctic ice core records (9)

**Figure 2.**
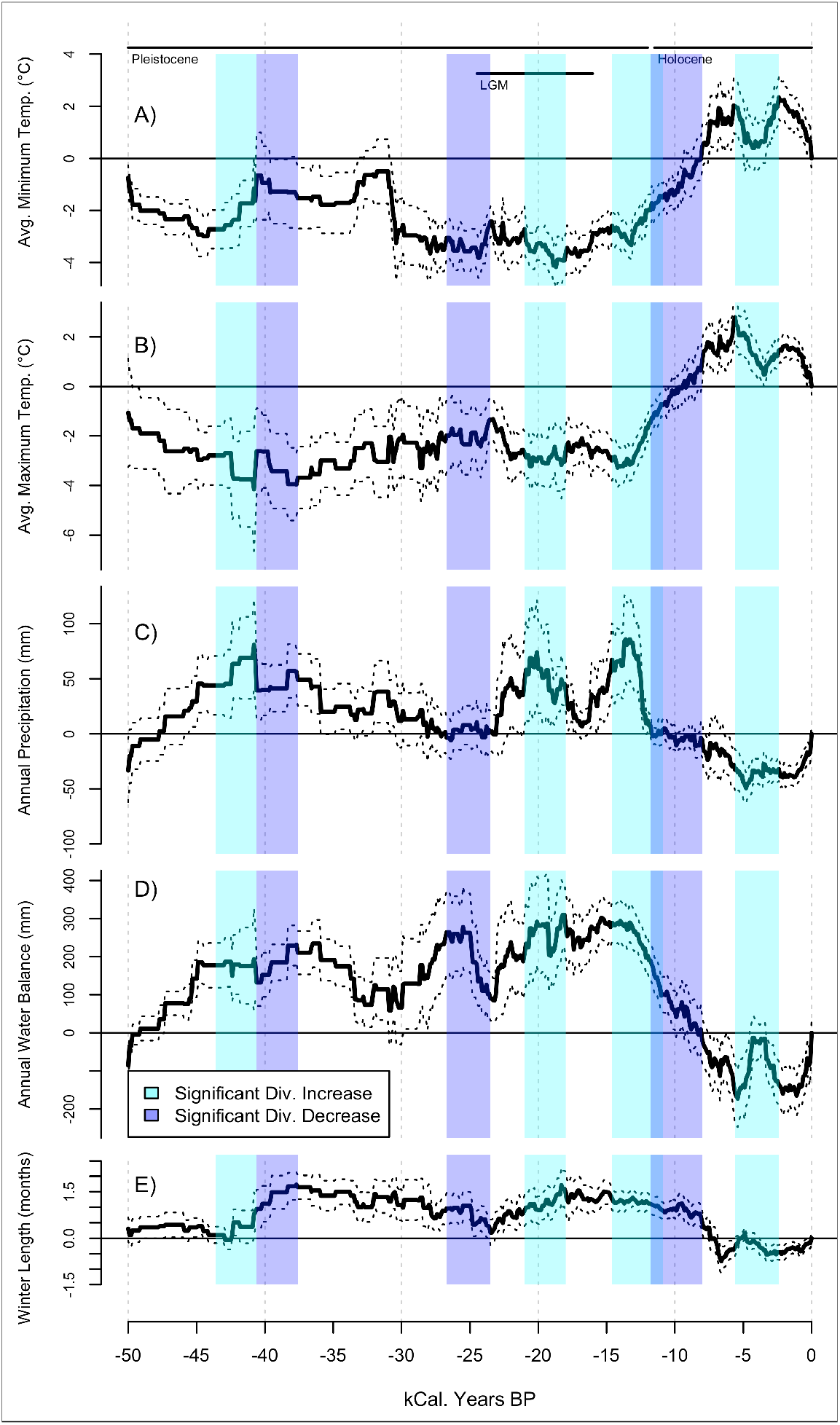
Temperature extremes, precipitation, water balance, and winter season CRACLE anomaly time series. A) Maximum annual temperature (°C), B) minimum annual temperature (°C), C) mean annual precipitation (mm), D) mean annual water balance (mm) – potential evapotranspiration + precipitation, E) estimated winter length (number of months with mean temperature less than 5°C). Solid lines indicate mean difference, dashed lines are +/− 2*SE. Shaded regions indicate time periods where significant increases (blue) or decreases (purple) in taxonomic diversity are observed (Supplementary Figure S1).

**Figure 3.**
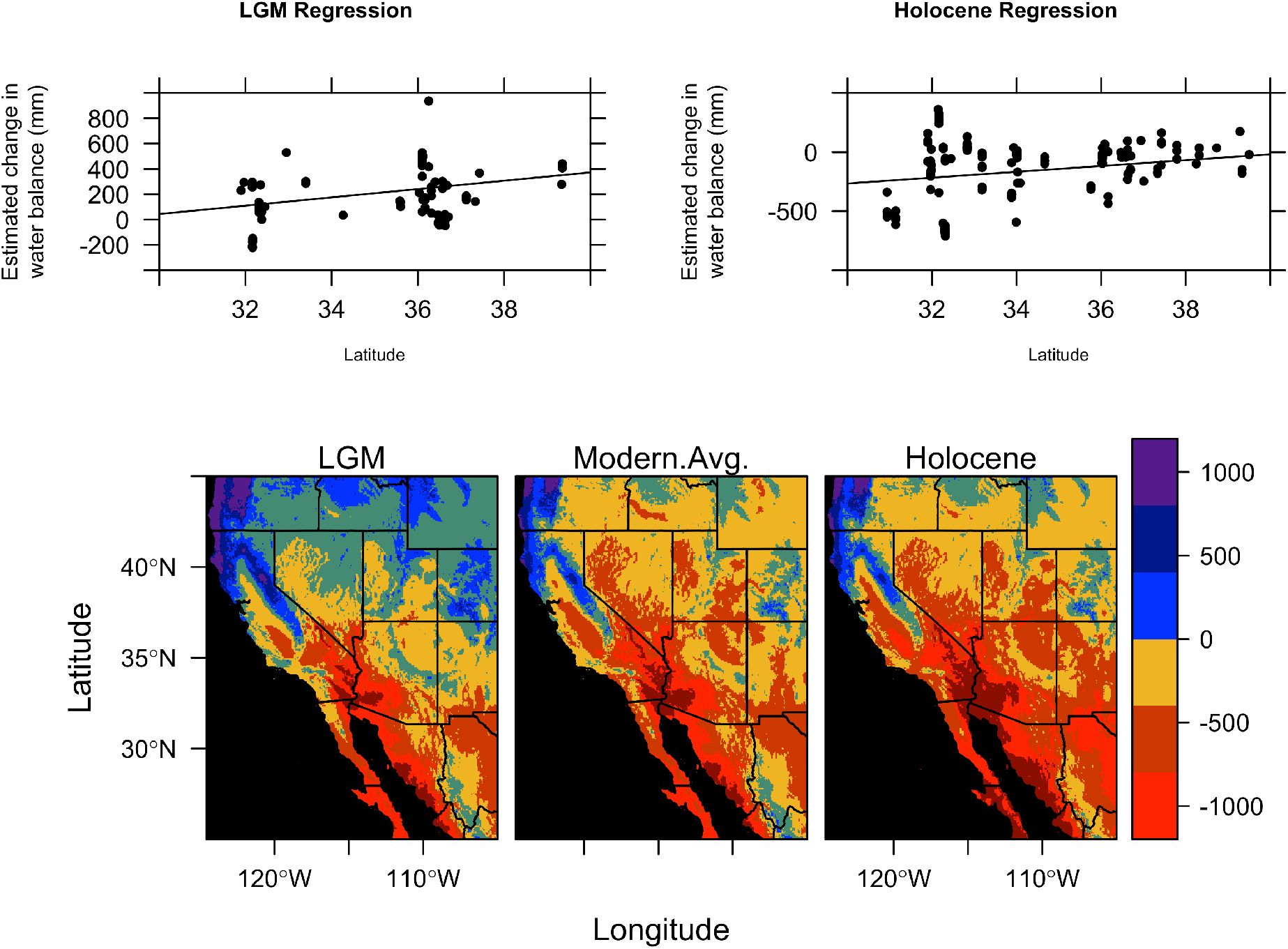
Linear modeling of Great Basin water balance (PET + MAP) during the estimated periods of greatest temperature departure from modern. A positive relationship exists between CRACLE inferred water balance anomalies vs. latitude for both the temperature minimum of the from 22kya to 15kya during the Last Glacial Maximum (*r*^2^=0.0927; *p*=0.004109; *n*=77) and the Holocene thermal optima between 8ky and 2kya (*r*^2^=0.1053; *p*=0.000; *n*=176) exhibit a positive and near-linear relationship. Applying the linear models to predict the water balance across the study region yields continuous adjusted estimates of water balance for the LGM and Holocene and visual comparison to the modern (1950-2000) average (77). Maps were generated with R version 3.3.2 (76) and associated geospatial libraries *raster* version 2.5-8 (82) and *rasterVis* version 0.41 (83).

## ACKNOWLEDGEMENTS

The authors would like to thank many colleagues for generous feedback on this manuscript through many drafts, especially Dr. Maria Alejandra Gandolfo-Nixon, Dr. William Crepet, Camilla Martinez, Thereis Choo, and Daniella Allevato. Funding support for this study was provided, in part, by the generous support of the Gerstner Family Foundation and the American Museum of Natural History.

## ADDITIONAL INFORMATION

### Author contributions

RSH and KCN contributed equally to this manuscript and the conception and design of this study. CRACLE Software was written and implemented by RSH. Analyses and synthesis were conducted by RSH.

